# Human RAD52 stimulates the RAD51-mediated homology search

**DOI:** 10.1101/2022.06.27.497340

**Authors:** Ali Akbar Muhammad, Thibault Peterlini, José Guirouilh-Barbat, Xavier Veaute, Didier Busso, Bernard Lopez, Jean-Yves Masson, Gérard Mazon, Eric Le Cam, Pauline Dupaigne

## Abstract

Homologous recombination (HR) is a DNA repair mechanism of double strand breaks and blocked replication forks that involves a process of homology search and homologous pairing leading to the formation of synaptic intermediates whose architecture and dynamics are tightly regulated to ensure genome integrity. In this mechanism, RAD51 recombinase plays a central role and is supported by many partner including BRCA2 and RAD52. If the mediator function of BRCA2 to load RAD51 on RPA-covered ssDNA is well established, the role of RAD52 in HR, more precisely its interplay with BRCA2 is still far from understood. We have used Transmission Electron Microscopy combined with biochemistry to characterize the sequential participation of RPA, RAD52 and BRCA2 in the assembly of the RAD51 filament, its architecture and its activity. Despite our results confirm that RAD52 lacks a mediator activity, we observed that RAD52 can tightly bind to RPA-coated ssDNA, inhibit the mediator role of BRCA2 and form shorter RAD52- and RAD51-containing mixed filaments that are more efficient in subsequent homology search and formation of synaptic complexes and D-loops, resulting in more frequent multi-invasions as well. Through the characterization of the behavior of RAD52 and BRCA2, these results provide new molecular insights on the formation and regulation of presynaptic and synaptic intermediates during human HR.

## Introduction

Homologous recombination (HR) is an evolutionary conserved process that plays a pivotal role in genome stability, diversity and plasticity. HR is indeed a key repair pathway able to faithfully repair DNA damages including double-strand breaks (DSB) and DNA gaps by copying the error-free information from the template DNA normally present in the sister chromatid [1–3]. Defects in HR are associated with genetic instability, chromosomal aberrations, carcinogenesis and cell death [4]. The molecular mechanisms of HR can be dissected in a serie of “mechanistic steps” resulting in the gradual formation of reversible DNA intermediates. it is initiated by the formation of single-stranded DNA (ssDNA) through the resection of double stranded DNA (dsDNA) from a DSB end or enlargement of a ssDNA gap. The ssDNA generated by resection is initially covered by Replication protein A (RPA), and with the help of a number of protein mediators, the recombinase RAD51 can form by displacing RPA from this ssDNA a presynaptic filament able to search and pair with the homolog dsDNA donor giving rise to the formation of joint molecules known as synaptic intermediates. The D-loop is the stable joint molecule formed upon invasion of the homologous dsDNA donor by the presynaptic RAD51 nucleofilament after alignment of the complementary strands and subsequent displacement of the third strand. The invading strand then serves as a primer to start synthesis within the D-loop enabling the recovery of the information lost at the original break point. In the postsynaptic steps the different synaptic intermediates would be resolved through alternative sub-pathways involving multiple helicases and structure-selective nucleases [5]. As a result, most of the repair outcomes will be generated without crossing-over of the DNA molecules involved, while some crossover products may form by nucleolytic resolution of these synaptic intermediates enabling the recovery of the information lost at the original break point.

The assembly and regulation of the RAD51 filament on DNA are crucial for the proper formation of synaptic intermediates and their outcome. It is also now well established that RAD51 and some parners play additionnal roles in the protection of DNA from nuclease attack and extensive resection, at DSBs and during replication [6][7]. As a recombinase, RAD51 is an ATP-modulated protein that form right-handed helical filaments on DNA (mostly ssDNA) [8] in which the DNA is stretched non-uniformly by 150% with a gap every 3 nucleotides, each triplet following the B-shape of DNA [9]. Filament formation is a two steps mechanism: RAD51 nucleation on RPA-covered ssDNA then elongation by cooperative polymerization along DNA. HR Homology search and strand exchange processes rely on the remarkable structure and properties of this filament. RAD51 has two DNA binding sites, site I oriented inside the filament binds to ssDNA and site II allows to transiently contact dsDNA donor. The filament likely facilitates base-flipping of triplet units facilitating homology probing and recognition by triplet base increments [10–15]. The homology probing has been shown to be based on tracts of 8-nucleotide microhomology and transient interactions between the stretched single-stranded DNA within filament and bases in a locally melted or stretched DNA duplex [14–20]. The interaction between the nucleoprotein filament on ssDNA and the duplex DNA donor results in their incorporation into a three-stranded intermediate, the synaptic complex (SC), also known as a paranemic joint [21–23]. Two types of SC have been described, those in which DNA strand pairing is maintained by RAD51 (sensitive to deproteinization), and those in which the invading ssDNA of the filament and the complementary strand of the dsDNA donor are aligned and intertwined to form the new heteroduplex (resistant to deproteinization) [24,25]. In the latter case, the heteroduplex and the displaced strand form the Displacement loop (D-loop), an important HR intermediate required to prime DNA synthesis by the 3’ OH of the invading strand in the heteroduplex [26]. Many studies have contributed to the better understanding of homology search and the D-loop dynamics; however, the mechanistic steps the specific roles of associated RAD51-partners leading to RAD51-mediated SC are incompletely characterized.

In humans, many RAD51 partners have been identified playing roles in the filament formation, its architecture, but also its activity in searching for homology and the handling of the subsequent D-loop. RAD51 mediators are proteins helping filament assembly and stabilization either by accelerating RAD51 nucleation on RPA-ssDNA or by decelerating its dissociation from ssDNA. BRCA2 mediates the nucleation of RAD51 filaments onto ssDNA covered by RPA [27,28] whereas some RAD51 paralogs bind and remodel the presynaptic filament to a stabilized and flexible conformation [29]. BRCA2 also directly binds RAD51 through BRC repeats (in sub-stoechiometric conditions) and selectively targets RAD51 to ssDNA then reducing non-productive interactions with dsDNA [30–33]. A BRC peptide was also shown to intercalate between RAD51 protomers within the filament, inhibiting RAD51 ATPase activity, and thereby suppressing RAD51 release from DNA [34]. Finally, a postsynaptic function of BRCA2 has been proposed involving inhibition of RAD51 excess-mediated D-loop dissociation, highlighting a role for homeostasis between RAD51 and BRCA2 as an important factor of HR in mammalian cells [35,36]. In yeast saccharomyces cerevisiae, Rad52 is identified as the main HR mediator through Rad51 filaments nucleation catalysis [37–39] and has been shown to directly intercalate into the presynaptic filaments by forming with Rad51 mixed filaments rendering them more resistant to Srs2 antirecombinase activity [40]. Unlike its yeast homologue, human RAD52 was considered to be a dispensable activity in light of the lack of strong phenotypes for the RAD52 mutants in vertebrates [41,42] but it appears to play a role in HR and its deficiency is now known to be synthetically lethal with several key recombination proteins, notably involved in RAD51 filament formation, including BRCA2, and RAD51 paralogs [43,44]. Yet RAD52 shares a number of biochemical properties with its yeast counterpart, including the formation of ring oligomers [45–47], the ability to catalyze the annealing of complementary strands and its potential cooperation with RAD51 in strand exchange activity [48–50], but a mediator activity has never been identified. In addition, RAD52 was shown to have functions independent of RAD51 in alternative DSB repair pathways involving Single Strand Annealing (SSA) and Break Induced Replication (BIR) and in promoting DNA synthesis during replication stress [51,52]. The exact role of RAD52 in homologous recombination, particularly its potential participation in the early stages including RAD51 filament installation and homology search but also its interplay with BRCA2 are not yet well established. Because the complexity of the human HR mechanism is increased by the number of actors involved and their interplay, the timing and dynamics but also the stoechiometric equilibrium of these players have to be finely coordinated and regulated to prevent genome instability.

In this article, we have reconstituted the early steps of human HR with purified proteins in a reaction using a synthetic long DNA overhang substrate mimicking the DNA substrates that result after resection, with its dsDNA and ssDNA regions. We have observed by using Transmission Electron Microscopy (TEM) the sequential recruitement to the DNA substrate of RPA, RAD52 and BRCA2 as well as RAD51 in order to better understand the interplay between these different actors in the assembly of the RAD51 filament, its architecture and its activity. While our results confirm that Rad52 lacks a mediator activity, despite its ability to tightly bind RPA-coated ssDNA, we observed that RAD52 can inhibit the mediator role of BRCA2 and form shorter RAD52- and RAD51-containing mixed filaments that are more efficient in subsequent homology search and formation of synaptic complexes and D-loops, resulting in more frequent multi-invasions as well.

## Material and methods

### Cells

RG37 cell line was derived from SV40-transformed GM639 human fibroblasts and contained the DR-GFP HR reporter [53]. Cells were cultured in DMEM (Gibco) supplemented with 10% FBS.

### HR assay

50,000 cells were seeded 1 day before siRNA transfection, which was carried out using INTERFERin following the manufacturer’s instructions (Polyplus Transfection) and 40 pmol siRNA: RAD52 (cat# AM16708) was purchased from Ambion and Control (5’- AUGAACGUGAAUUGCUCAA −3’) and siRAD51 (5’-GUGCUGCAGCCUAAUGAGA-3’) were synthesized by Eurofins. Forty-eight hours later, cells were transfected with the pBASce-HA-I-SceI expression plasmid with Jet-PEI following the manufacturer’s instructions (Polyplus Transfection). Cells were detached 72 hours after I-SceI transfection and analyzed by flow cytometry for the expression of the GFP reporter.

### Western blot analysis

Cells were lysed 30 minutes at RT in a buffer containing benzonase (>250 U/mL) in 50 mM Tris-HCl (pH 7.5), 20 mM NaCl, 1mM MgCl_2_, 0.1% SDS, supplemented with a complete mini protease inhibitor (Roche). Proteins (40 μg) were denatured 10min at 55°C, electrophoresed on 9% SDS-PAGE gels, transferred onto nitrocellulose membranes and probed with the following specific antibodies: anti RAD52 (sc-365341, Santa Cruz) anti-RAD51 (PC130, Millipore), and anti-vinculin (Abcam). Immunoreactivity was visualized using an enhanced chemiluminescence detection kit (ECL, Pierce).

### Protein purification

Human RAD51 was purified on CiGEX Platform (CEA, Fontenay-aux-Roses) as following. His-SUMO-RAD51 was expressed in *E. coli* strain BRL (DE3) pLys. All of the protein purification steps were carried out at 4°C. Cells from a 3-liter culture that was induced with 0,5 mM isopropyl-1-thio-ß-D-galactopyranoside for overnight at 20°C were resuspended in PBS x1, 350 mM NaCl, 20 mM Imidazole, 10% Glycerol, 0,5 mg/ml Lysozyme, Compete Protease Inhibitor (Roche), 1 mM 4-(2-aminoethyl)benzenesulfonyl fluoride (AEBSF). Cells were lysed by sonication and the insoluble material was removed by centrifugation at 150,000 x g for 1h. The supernatant was incubated with 5 ml of Ni-NTA resin (Qiagen) for 2h. The mixture was poured into an Econo-Column Chromatography Column (BIO-RAD) and the beads were washed first with 80 ml W1 buffer (20 mM Tris HCl pH 8, 500 mM NaCl, 20 mM Imidazole, 10% glycerol, 0,5% NP40), followed by 80 ml of W2 buffer (20mM Tris HCl pH 8, 100mM NaCl, 20mM Imidazole, 10% glycerol, 1 mM DTT). his-SUMO-RAD51 bound to the beads was then resuspended with 8ml of W2 buffer and incubated with SUMO protease at a ratio 1/80 (W/W) for 16 h. RAD51 without the his-SUMO tag was then recovered into the flow thru and directly loaded onto a HiTrap heparin column (GE Healthcare). The column was washed with W2 buffer and then a 0.1-1M NaCl gradient was applied. Fractions containing purified RAD51 were concentrated and dialyzed against storage buffer (20mM Tris HCl pH 8, 50mM KCl, 0.5 mM EDTA, 10% glycerol, 1 mM DTT, 0.5 mM AEBSF) and stored at −80°C.

Human RPA protein was purified on CiGEX Platform (CEA, Fontenay-aux-Roses) as previously described [54]. RAD52 and BRCA2 were purified as previously described [55].

### Synthesis of 3’ overhang DNA construction (400 base pairs with a 3’ overhang of 1040 nucleotides)

Two DNA fragments of 1040 and 400 bp were amplified from pBR322 plasmid by PCR using *Taq* polymerase and the pairs of primers Cy5-2574^+^ x biotin-4014^-^ and biotin-2574^+^ x 2976^-^, respectively. The biotinylated PCR products were purified on a MiniQ 4.6/50 ion exchange column (GE Healthcare Life Sciences) and loaded into a HiTrap Streptavidin HP column (Amersham Biosciences). Purification of the non-biotinylated strand was achieved by elution with 80 mM NaOH, neutralized by addition of HCl 1M and annealed at equimolar concentrations in molecules, in presence of 1.5 mM MgCl_2_ then purified on an ion exchange MiniQ column.

### DNA-protein complexes for TEM statistical analysis

For RPA-DNA complexes and mediation assay, the ssDNA part of the 3’ overhang DNA substrate was covered with saturated concentration of RPA as following. 15 μM in nucleotides of the DNA substrate were incubated with 0,3 μM RPA (1 protein per 20 nt of ssDNA) in a buffer containing 10 mM Tris-HCl pH7.5, 50 mM NaCl 10 minutes at 37°C. BRCA2 (2 to 5 nM) and/or RAD52 (0,1 to 0,5 μM) were then introduced in the reaction during 15 minutes at 37°C, and for the mediation assays, 5 μM RAD51 were added at the same time as BRCA2/RAD52 partner. In the last experiment, the buffer was completed with 2 mM MgCl_2_, 2 mM CaCl_2_, 1.5 mM ATP and 1 mM DTT.

RAD51 filaments were formed by incubating 15 μM in nucleotides of 3’ overhang DNA labeled with Cy5 with 5 μM RAD51 (1 protein per 3 nt) in a buffer containing 10 mM Tris-HCl pH7.5, 50 mM NaCl, 2 mM MgCl_2_, 2 mM CaCl_2_, 1.5 mM ATP and 1 mM DTT 3 minutes at 37°C, then adding 0.25 μM RPA (1 protein per 60 nt) during 15 minutes. BRCA2 (2 to 5 nM) or RAD52 (0,1 to 0,5 μM) was added in the reaction at the same time as RAD51.

### Transmission Electron Microscopy

For DNA-protein complexes observation, 2 μL of the remaining 7 μL of reaction were quickly diluted 120 times in a buffer containing 10 mM Tris-HCl pH 7.5, 50 mM NaCl, 2 mM MgCl_2_, 2 mM Cacl_2_ and observed by electron microcopy (DNA-protein samples). During one minute, a 5 μL drop of the dilution was deposited on a 600-mesh copper grid previously covered with a thin carbon film and pre-activated by glow-discharge in the presence of amylamine (Sigma-Aldrich, France) [56,57]. Grids were rinsed and positively stained with aqueous 2 % (w/v) uranyl acetate, dried carefully with a filter paper and observed in the annular dark-field mode in zero loss filtered imaging, using a Zeiss 902 transmission electron microscope. Images were captured at a magnification of 85,000× with a MegaviewIII Veleta CCD camera and analyzed with iTEM software (both Olympus Soft Imaging Solution). For the quantifications, the different populations of molecules were counted on at least 2 independent experiments with a total of at least 200 molecules counted.

### D-loop *in vitro* assay and analysis of the DNA-protein and DNA intermermediates

In the first step of the reaction the RAD51 filament was assembled as following. 15 μM in nucleotides of 3’ overhang DNA labeled with Cy5 were incubated with 5 μM RAD51 (1 protein per 3 nt) in a buffer containing 10 mM Tris-HCl pH7.5, 50 mM NaCl, 2 mM MgCl_2_, 2 mM CaCl_2_, 1.5 mM ATP and 1 mM DTT 3 minutes at 37°C. Then 0.25 μM RPA (1 protein per 60 nt) were added during 15 minutes. In the second step, 25 nM in molecules of homologous/heterologous dsDNA donor was introduced during 30 minutes at 37°C. For the homologous donor, pUC19 plasmid was used, while PhiX174 RFI was used as heterologous DNA, both purchased from New England Biolabs and purified on MiniQ ion exchange chromatography column.

### D-loop assay analysis by agarose gel electrophoresis

7 μL of the 14 μL D-loop reaction reaction were taken and stopped using 0.5 mg/mL Proteinase K, 1% SDS, 12.5 mM EDTA and incubated overnight at room temperature. A 1% TAE agarose gel was run at 70 V, for 40 minutes.

### RAD52 immunolabeling for its TEM detection

To test the presence of RAD52 in the RAD51 filament and in joint molecule structures, we carried out an immunoaffinity labeling procedure. The DNA-protein complexes formed in 7 μL of reaction were first stabilized using incubation with 0.01 % glutaraldehyde cross-linking agent during 10 minutes at 30°C. Then 3 μM of a polyclonal anti yeast RAD52 antibody (primary antibody rabbit IgGs) were added to the reaction and incubated at 25 °C during 10 minutes followed by 10 minutes incubation with 5 μM of the secondary immunogold antibody (anti rabbit). The labeled reaction was then crosslinked with 0.04 % glutaraldehyde (0.05 % in final concentration) for its subsequent purification by gel filtration on smart system using a superose 6 column (Amersham). This last procedure is essential to remove the excess of proteins and primary/secondary antibodies that are not bound to the complexes. To ensure that the labeling is specific to RAD52 detection, the controle experiments with pure RAD51 filaments (in absence of RAD52) were carried out. Samples were visualized by EM in dark fields and bright field modes. Note that although this procedure is useful for demonstrating the presence of a protein, the identification remains qualitative.

### Statistical analysis

Statistical analysis were performed with Prism 9 (GraphPad software). The statistical tests used are indicated in the legends of the figures. ns: not significant, *: P<0.05, **: P<0.01, ***: P<0.001, ****: P<0.0001.

## Results

### RAD52 is involved in HR in response to DSB

Different from its yeast homolog where Rad52 defines the epistasis group of proteins for HR pathways [58], mammalian RAD52 is not an essential protein for HR and RAD52 -/- mice are viable, fertile and only show a slight decrease in HR activity [41]. In general, deficiency of RAD52 is not linked to direct evidences of damage sensitivity but the fact that overexpression of RAD52 in mammalian cells improves their resistance to ionizing radiation suggests a role in the response to DNA damage by HR [59]. To further precise its involvement in HR, we decided to use a dedicated reporter system, the DR-GFP assay [53], able to monitor mild recombination phenotypes (see figure 1). This DR-GFP assay tracks the restoration of GFP expression as a result of an I-SceI-induced (DSB) gene conversion. Cellular green fluorescence is then assessed to quantify gene conversion frequency. While the silencing of recombinase RAD51 through siRNA totally abolishes HR gene conversion in the DR-GFP system, the RAD52 silencing leads to a partial decrease in gene conversion that dropped to 40% of its wild-type levels, clearly demonstrating the importance of RAD52 for DSB-induced gene conversion through HR (figure 1 C).

**figure 1:**
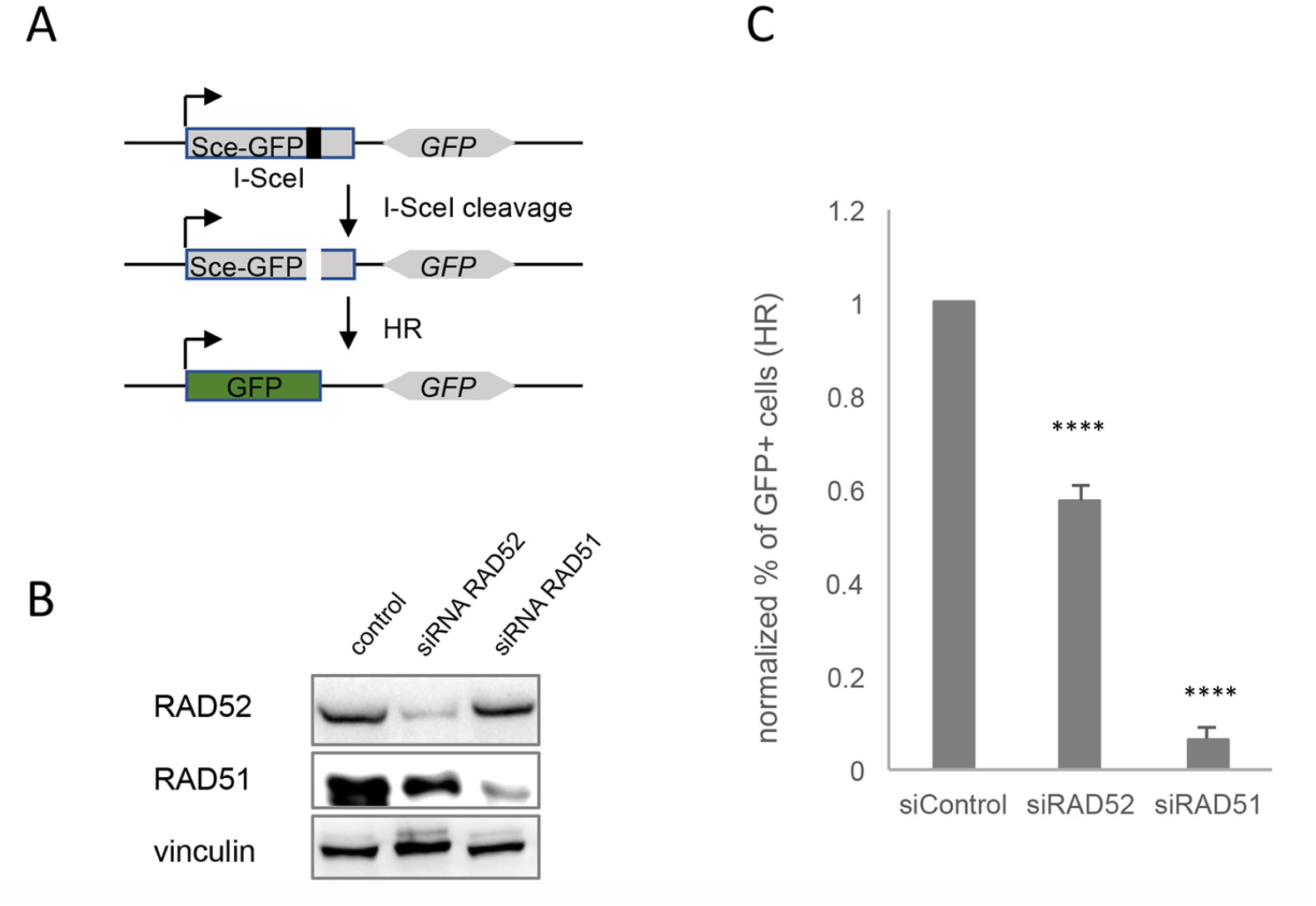
Homologous Recombination is affected in cells silenced for RAD52. A. Schema of the DR-GFP reporter assay used to monitor HR (PMID: 10541549) . SceGFP is a modified GFP gene containing an I-SceI site and in-frame termination codon. *GFP* is an 812-bp internal GFP fragment. Repair by HR (gene conversion) results in a functional GFP gene. B. Depletion of RAD51 and RAD52 in cells transfected with RAD51 and RAD52 siRNA. C. Normalized frequency of HR in cells silenced for RAD51 or RAD52. Bars are mean ±SEM, and reflect the results of 6 independent experiments. (****= p<0.0001, unpaired t-tests, two tailed).

### RAD52 binds and compacts RPA covered ssDNA

To further clarify the role of RAD52 in the HR mechanism, we implemented biochemical reactions and first tested its ability to bind RPA covered ssDNA on a ss-dsDNA hybrid substrate mimicking the ssDNA overhang generated after a DSB resected end (figure 2). The substrate contained a dsDNA region of 400 bp and 3’ overhang of 1040 nucleotides to mimic the structure and approximate length found in DSBs processed *in vivo* (figure 2 A) [60]. After incubation of this 3’ overhang substrate with saturating concentrations of RPA we observed its binding covering the ssDNA part (figure 2 B). RPA was bound without a defined structure, thus reflecting the flexible and stochastic nature of RPA binding to ssDNA. As can be observed in figure 2B, addition of RPA allowed the deployment of the ssDNA on the surface of the TEM grid by destabilizing secondary structures (figure 2 B compared to 2 A - naked DNA substrate), while the dsDNA part of the substrate remains linear and well spread on the grid surface. The addition of RAD52 to the reactions containing RPA complexed with the 3’ overhang substrate first revealed that RAD52 can bind to RPA-ssDNA and form discrete complexes (figure 2 C, D) accompanied by a significant decrease in the RPA-ssDNA length of nearly 30 % of the size before RAD52 addition (from mean length 314+/-77 nm to 220+/-93 nm, figure 2 E). RAD52 is known to oligomerize adopting a ring shape [45–47,61], such oligomerization could explain this shortening as the RPA-ssDNA fibers folds or winds around RAD52 oligomers. While we did not observe any interaction of RAD52 with dsDNA part, we signaled a very frequent localization of RAD52 complexes at the dsDNA-ssDNA junction (37 % +/- 7), which could be explained by a sliding of RAD52 along the RPA-sbDNA to this junction (supplementary figure S1 A-C). RAD52 could also interact tightly with the naked ssDNA of our DNA substrate but in this conditions it did not form discrete complexes as those observed in the presence of RPA but it rather showed a tendency to form aggregates and DNA intramolecular bridges (Supplementary Figure S2 A, B), as previously observed [62]. Interestingly, we showed that a 4 nucleotides long overhang was sufficient to promote RAD52 recruitment at the ss-ds DNA junction with the formation of discrete RAD52-DNA substrate complexes coexisting with end to end aggregated DNA substrates (Supplementary Figure S2 C-E). This aggregation of DNA molecules was also obtained when increasing the RAD52 concentration, revealing an intrinsic property of RAD52 to gather and hold together ssDNA molecules, maybe in relation to its ability to anneal complementary DNA sequences. These properties of RAD52 can be explained by the presence in the RAD52 oligomeric ring of two DNA-binding sites, a primary ssDNA-binding site located at the circumference of the ring and a second DNA-binding site that can bind either ssDNA or dsDNA [45,47,61,63,64].

**figure 2:**
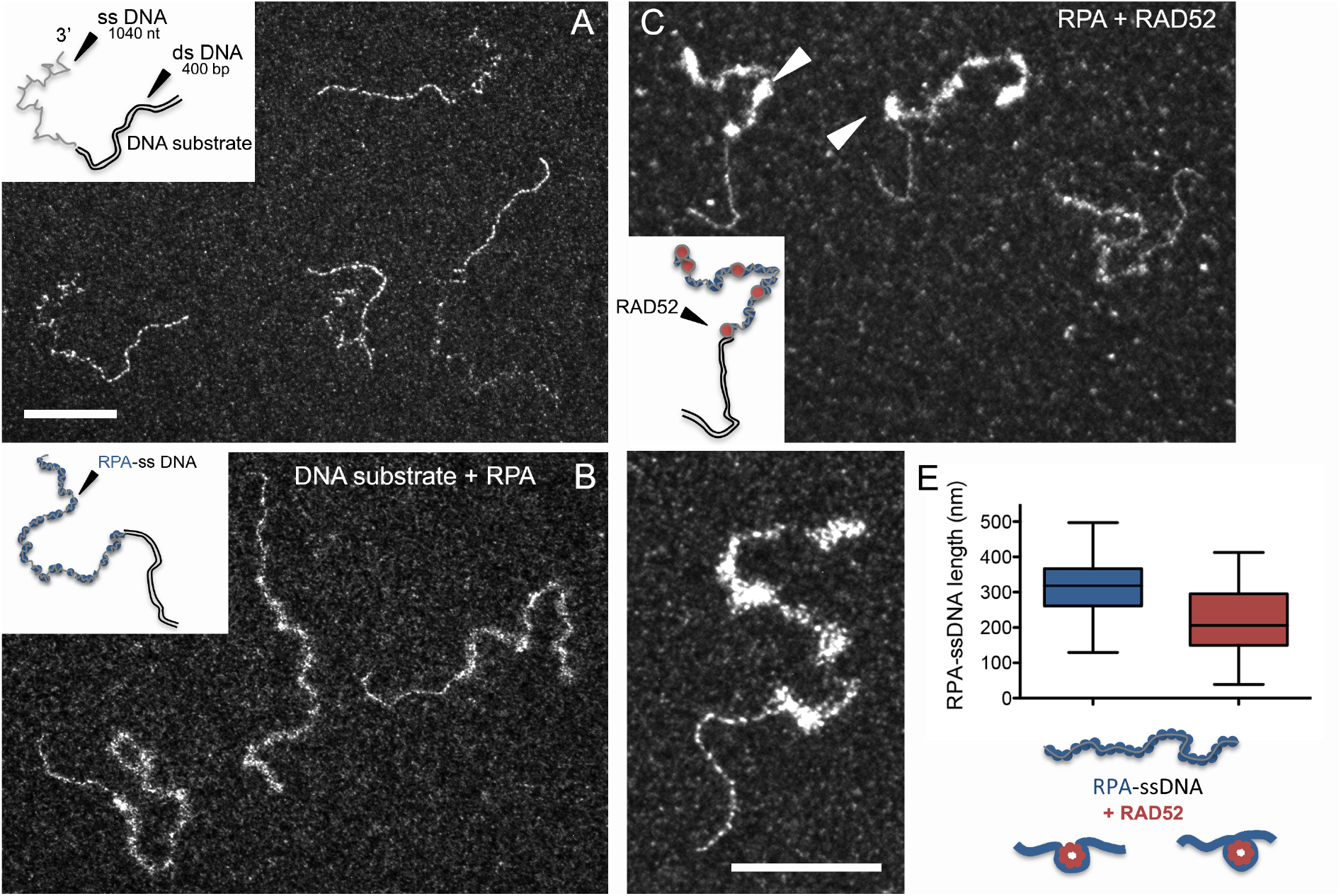
RAD52 binds to RPA-ssDNA. (A-D). Representating TEM images of the DNA-protein complexes in the reactions with insets of schematic drawings of the molecules. A. DNA substrate containing a 400 bp dsDNA prolongated with a 1040 nt long ssDNA overhang. B. The substrate is covered by saturated concentration of RPA protein. C. Then RAD52 is added in the reaction, discrte complexes are detected and pointed with arrows. (A-D: same magnification, the scale bar represent 100 nm). D. zoom on a RPA-RAD52-DNA complex (the scale bar represent 100 nm). E. Measurements of the length of RPA-ssDNA in absence (in blue) or in presence (in red) of RAD52. RAD52 induces a 30% length reduction.

### RAD52 bound to RPA-ssDNA inhibits BRCA2 mediator activity

We then decided to test the putative mediator activity of RAD52 and BRCA2 individually or combined by analyzing their capacity to recruit RAD51 on the RPA-covered DNA overhang substrate thus resulting in RAD51-filament formation (figure 3). Precisely, the DNA substrate was first incubated with saturating amounts of RPA to generate RPA-covered ssDNA overhangs, and RAD51 was then added either in the presence of purified RAD52 and/or BRCA2 proteins. The presence of pre-bound RPA clearly inhibited RAD51 loading and filament assembly on ssDNA part (figure 3 A), which was consistent with the fact that *in vitro* formation of RAD51 filaments on ssDNA usually requires RPA to be added to the reaction mixture after RAD51. Interestingly, in these conditions, RAD51 was able to polymerize on the dsDNA section of the substrate forming continuous filaments, the ssDNA still being covered by RPA (figure 3 A, E). It is worth noting that in the absence of RPA, we found that human RAD51 similarly binds to ssDNA or dsDNA (Supplementary Figure S3 A), in contrast to *Saccharomyces cerevisiae’s* Rad51 or *Escherichia coli’s* RecA recombinases, which exhibit a preferencial affinity for ssDNA. This specific property of hRAD51 raises the question of whether there is a transitory binding of RAD51 to dsDNA that should be negatively regulated, as observed in some contexts *in vivo* [65].

**figure 3:**
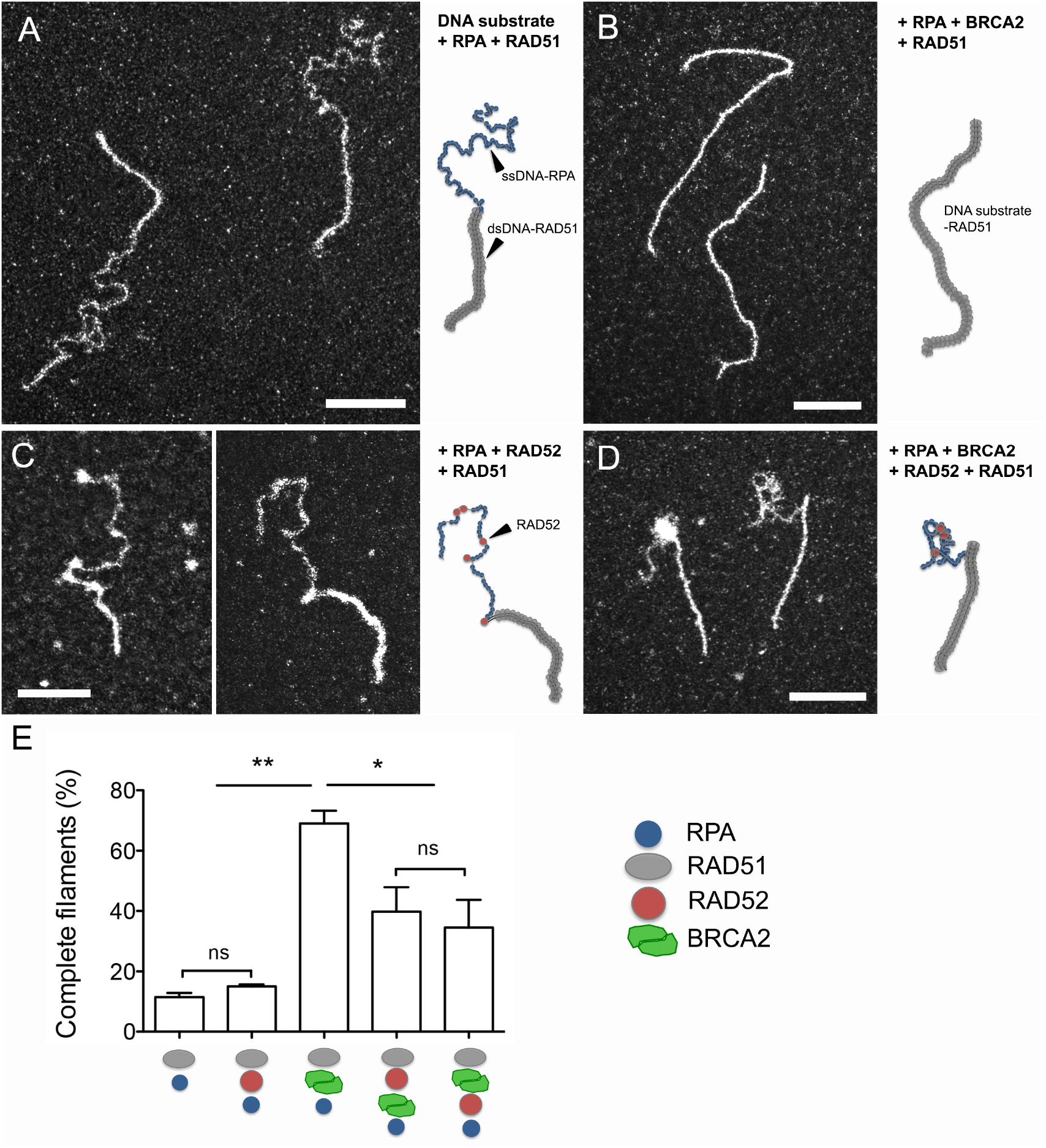
early introduction of RAD52 inhibits RAD51 assembly on ssDNA. (A-D). Representating TEM images of the DNA-protein complexes in the reactions with insets of schematic drawings of the molecules. A. DNA substrate is first incubated with staurated concentration of RPA then RAD51 (1 protein per 3 nucleotides) is added in the reaction. RAD51 filament assembles on the dsDNA part of the substrate without replacing RPA on the ssDNA part. B. sub-stoechiometric amount of BRCA2 (2 nM) is introduced in the reaction simultaneously with RAD51 and promote the formation of complete filaments (on dsDNA but also ssDNA parts of the substrate) then replacing RPA along ssDNA, highlighting BRCA2 mediator role. C. 0,2 μM RAD52 were added in the reaction simultaneously with RAD51. RAD52 binds to RPA-ssDNA but does not allow RAD51 filament installation on ssDNA confirming that human RAD52 does not exhibit any mediator activity as BRCA2. D. RAD52 and BRCA2 are introduced together in the reaction. RAD52 binding to RPA-ssDNA predominantly prevents the BRCA2-mediated nucleation of RAD51 on ssDNA and subsequent complete filament formation, then partially inhibiting BRCA2 mediator activity of BRCA2. All scale barsreprsent 200 nm. E. Quantification of complete RAD51 filaments (assembled on dsDNA and ssDNA parts of the DNA substrate) in the DNA-protein samples. Error bars indicate standard deviation from three independent experiments.

The introduction of BRCA2 at sub-stoechiometric concentration (from 1 to 5 nM - one protein for one DNA substrate) promoted the loading and assembly of RAD51 by replacing RPA on the ssDNA part of the overhang to form complete filaments on the whole substrate (figure 3 B). This bona fide mediator activity was illustrated by the presence of 69% complete RAD51-filaments in the presence of 1 nM BRCA2 compared to 10% in its absence (figure 3 E). This mediator activity of BRCA2 couldn’t be substituted by adding RAD52 to the reaction (at concentrations ranging from 0,1 to 1 μM), which did not allow RAD51 loading on RPA-covered ssDNA (figure 3 C, E). Again, RAD52 discrete fixation on RPA-ssDNA was detected, coupled to a significant reduction of the length of RPA-ssDNA complexes. When both BRCA2 and RAD52 were introduced together in the reaction, regardless of the order they were added, a partial inhibition of the BRCA2 mediator activity was observed with roughly a 50% decrease of the complete filaments that could be observed (figure 3 D-E). This reduction is compatible with a competition between RAD52 and RAD51-BRCA2 for the binding to RPA-ssDNA.

### RAD52 participates in the formation of mixed fragmentized filaments whereas BRCA2 catalyzes formation of long and continuous RAD51 filaments

To further analyze the effect of RAD52 and/or BRCA2 on the architecture and activity of RAD51 filaments, we carried out a series of reactions where we first incubated the DNA overhang substrate with RAD51 and either RAD52 or BRCA2, followed by addition of a non-saturating amount of RPA to help in the filament installation through removal of ssDNA secondary structures upon RAD51 polymerization (figure 4).

**figure 4:**
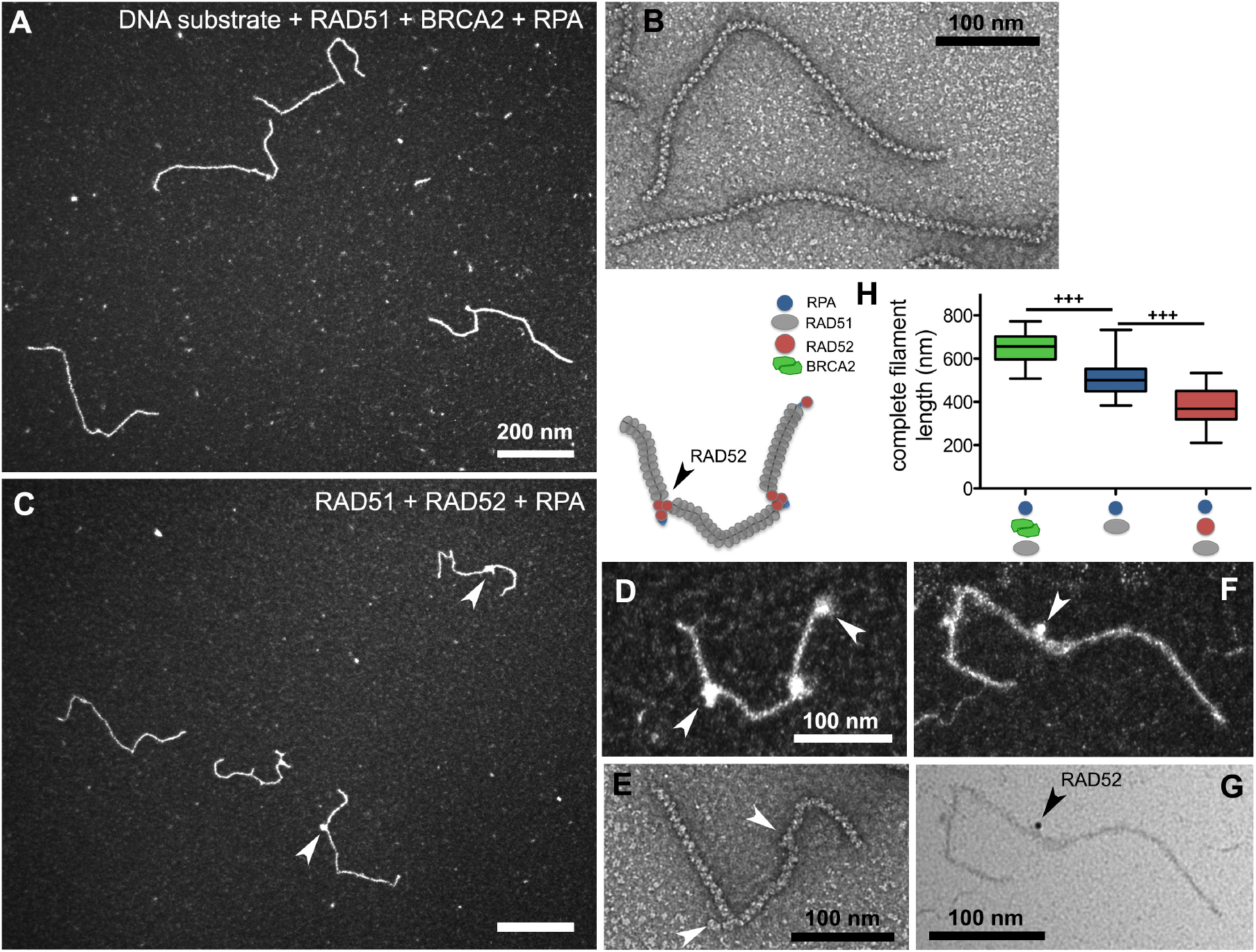
BRCA2 induces the formation of long continue RAD51 filaments while RAD52 forms with RAD51 mixed and fragmented filaments. A. Representative TEM image of RAD51 filaments formed in presence of 5 nM BRCA2. B. Negative staining TEM image showing the same long and continue filments. C. Mixed filaments formed in presence of 0,2 μM RAD52. Arrows point bright spots in the filaments suggesting RAD52 discrete complexes bound to the filment. D. Zoom on a mixed filament with schematic drawing of the molecule above. E. Negative staining image of a mixed filament. F and G. Darkfield and brightfield TEM imaging of Immunostaining experiment of mixed RAD51-RAD52 filament specifically showing the presence and localization of RAD52 using anti-RAD52 antibody and gold beads coupled secondary antibody. Arrow point the gold bead (with increased electron density). H. Measurement of the length of complete filaments formed in presence of BRCA2 (in green), pure filaments formed by RAD51 then low mount of RPA (in blue) and mixed RAD51-RAD52 filaments (in red).

The presence of BRCA2 (5nM) in the reaction promoted the formation of long, complete and continuous filaments, that thanks to a negative staining TEM observation were revealed to show a regular helical architecture without interruptions (figure 4 A, B and H). We couldn’t detect any BRCA2 binding onto RAD51 filaments, thus suggesting that BRCA2 may promote RAD51 recruitment without remaining stably bound to ssDNA in our experimental conditions. To note that BRCA2 is a 380 kDa protein forming a dimer in solution, its size would allow its detection within the RAD51 filament [66]. RAD51-filaments formed in presence of BRCA2 in these conditions were 27 % longer than those formed in its absence (648 +/-66 nm, which corresponds, after correction for DNA extension by RAD51 to 1270 +/-129 pb, figure 4 H) indicating that BRCA2 not only had a positive effect on RAD51 nucleation, but also on the filament elongation and stability. Moreover, the addition of BRCA2 on pre-assembled filaments also resulted in a significant increase in their length, further indicating a role of BRCA2 on filament stability. Different from what observed in the presence of BRCA2, RAD51-filaments formed in the presence of RAD52 (0,2 μM) displayed a number of discontinuities and the presence of discrete complexes in the form of 2 to 3 bright spots per filament, detectable using either a positive or a negative staining of the sample (figure 4 C-E). The presence of these clusters or complexes suggested RAD52 was part of the RAD51 filament and its helicity interruption was often associated with a ‘kink’ in the filament. To specifically show and localize RAD52 inside the filaments, we performed immunolabelling assays by using an anti-RAD52 antibody and a secondary antibody coupled to gold beads. This labeling enabled observing, indeed, how mixed RAD51-RAD52 filaments consist in tracts of RAD51 filament interrupted by RAD52 oligomers, specifically tagged by the presence of gold beads (figure 4 G, F). Increasing the RAD52 concentration above 1 μM RAD52 did not induce an increase in the number of discrete spots per filaments but rather caused a background covering with particles (supplementary figure S3 B) pointing to competition between RAD51 and RAD52 for ssDNA binding thus allowing only a limited number of oligomers to interact with the filament. Again, RAD52 binding to the RAD51-filament was associated with a significant 25% reduction of the filament length, in line with a putative winding of the ssDNA or RPA-ssDNA fiber around the RAD52 oligomers (figure 4 H). Addition of RAD52 in a later time point of the reaction, over an already pre-formed RAD51 filament, did not change the shape, length and helical architecture of the filament, indicating that RAD52 in this conditions was excluded from the filament, a result in agreement with Greene et al. 2017 [67].

### Human RAD51 displays an important ability to contact a dsDNA donor independently of the presence of sequence homology

It was previously shown that human RAD51 can form D-loops on its own, but only in the presence of Calcium, this activity being stimulated by RAD54, while its yeast homolog absolutely requires Rad54 to form synaptic complexes and D-loops [68–71]. We decided to test whether this strand invasion activity of RAD51 is influenced by the presence of BRCA2 or RAD52 (figure 5).

**figure 5:**
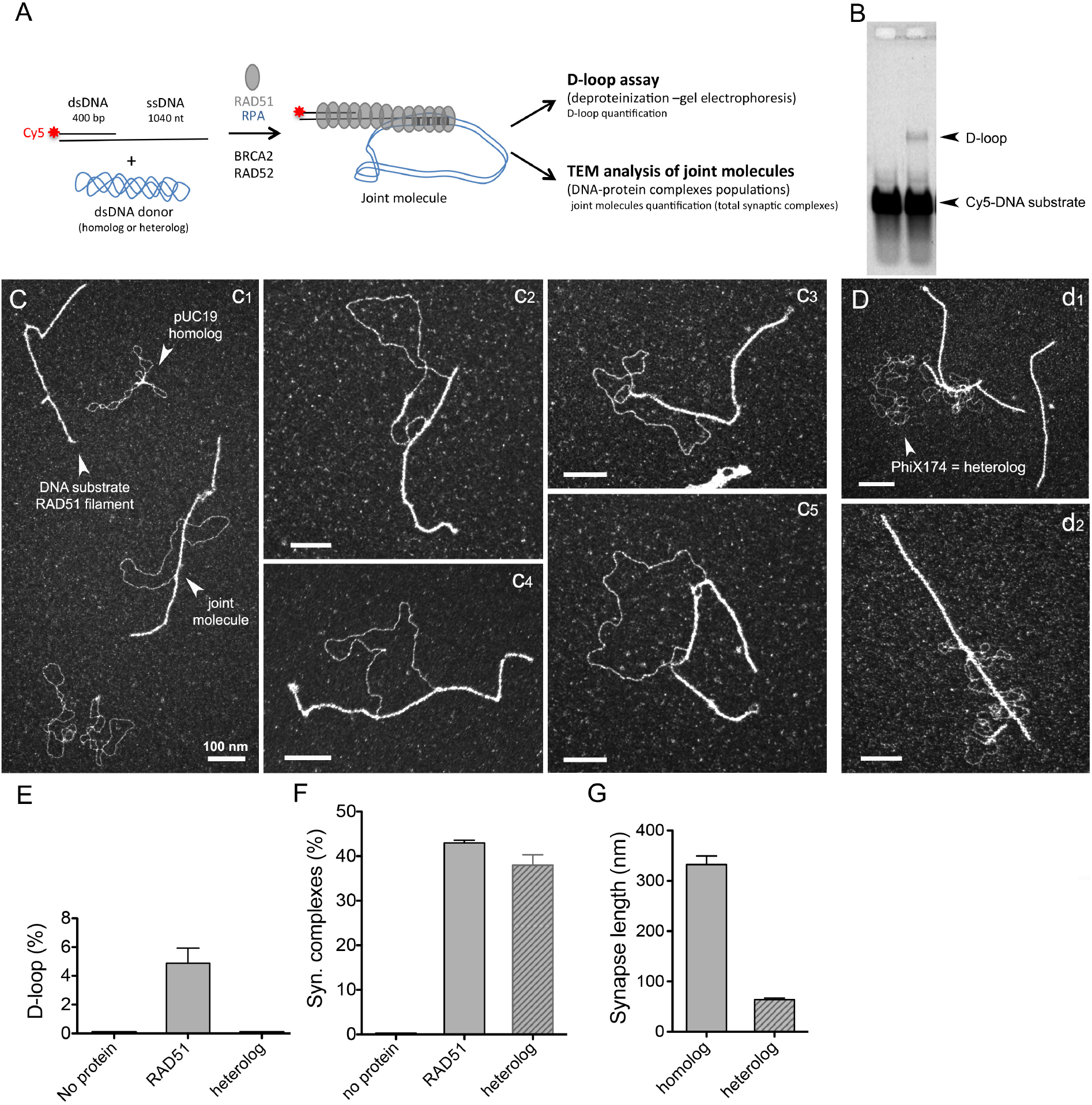
human RAD51 filament is highly active in contacting dsDNA donor to form synaptic complexes. A. Scheme of the D-loop *in vitro* reaction. The RAD51 filament is preformed on the ss-dsDNA substrate then dsDNA heterologous or homologous donor is added to the reaction. Three independant reactions have been performed. For each reaction, one part is deproteinized then run on a 1% agarose gel. D-loops are quantified using image J software. The second part is diluted, spread on a microscopy grid and analyzed upon TEM. B. representative gel of the D-loop assay. C-D. TEM images of the reaction performed using dsDNA donor homologous (C) or heterologous (D). The scale bars in all images represents 100 nm.E. Quantification of the D-loop yield (for 3 independant D-loop reactions). F. Quantification of synaptic complexes formed by the pairing of RAD51 filament with dsDNA donor. Synaptic complexes percentage represents the number of filaments paired with dsDNA donor divided by the total number of filaments (paired and non-paired). G. Measurement of the synapse length (in nm).

A RAD51 filament was first pre-formed on the DNA substrate, followed by the addition of a dsDNA donor (containing homologous or heterologous sequences) to the reaction (figure 5 A). Homology search and strand invasion processes are characterized by the formation of joint molecules where the presynaptic filament pairs with the homologous dsDNA donor [72]. In our experimental approach, half the sample was subjected to TEM analysis, while the other half was analyzed in a traditional D-loop assay involving deproteinization of the reaction products, DNA species separation on a gel and quantification (figure 5 A). The D-loop is defined as the joint molecule product of the incorporation of the invading strand into a homologous dsDNA donor, resulting in the disruption of its original base pairing replaced by a newly formed heteroduplex. In the D-loop assay, D-loop products are characterized by the intertwining of the invading strand with its complement in the donor, they are stable in the absence of proteins and then resistant to deproteinization. With our overhang substrate, in the presence of RAD51 and an homologous dsDNA donor, 4,9 % D-loops were observed at 20 minutes after the dsDNA donor addition in the reaction (figure 5 B, E). Joint molecules were also directly visualized using TEM (figure 5 C, D). TEM is ideal to directly observe different populations of DNA and protein:DNA complexes. In the case of the D-loop reaction, this allowed us studying the intermediates that precede strand intertwining, specifically protein-mediated pairings such as synaptic complexes where the invading DNA is not intertwined with the donor molecule, these joint molecules being susceptible to deproteinization. We clearly distinguished joint molecules synaptic complexes resulting of the interaction between the nucleoprotein filament and the duplex DNA donor, coexisting with the reaction substrates (free RAD51 filaments on 3’ substrate and supercoiled DNA, figure 5 C). Surprisingly, at 20 minutes after dsDNA donor addition, 43 % of synaptic intermediates were counted, revealing the extraordinary capacity of human RAD51 filament to establish stable contacts with dsDNA (figure 5 F). Only 9 % of synaptic contacts resulted in the formation of D-loops by complementary sequences alignement (resistant to deproteinization and quantified by gel), indicating that the vast majority of synaptic complexes observed by TEM are potentially three stranded intermediates maintained by RAD51. Observation by TEM also enables the characterization of the joint-molecule architecture, as we can precisely determine where the proteins are bound and thus measure the DNA length as well as the synaptic part of the joint molecules. Analyzing RAD51-mediated synaptic complexes (SC) we observed that the contact zone (synapse) between the filament and the dsDNA donor remained covered by RAD51 with an average contact length of 332 ±60 nm, equivalent to 651 bp when corrected for the extension by RAD51 (figure 5 G). The topology of the negatively supercoiled dsDNA donor is affected by joint molecule formation as it relaxes as a consequence of SC and subsequent D-loop formation [3,21,73,74]. The stretching of the donor in the SC by the RAD51 filament consumes negative supercoiling of the donor duplex. Interestingly when dsDNA donor added was heterologous, 38 % of SCs were observed (figure 5 D, F, G). Thus the formation of the major part of these stable RAD51-mediated contacts was independant of the presence of homology. However, qualitatively, joint molecules formed with heterologous dsDNA showed distinct features in comparison to the homologously-paired SCs. These nucleoprotein filament interactions with heterologous DNA were mainly characterized by short contacts (< 50 nm) and the dsDNA did not become topologically relaxed.

### RAD52 promotes synaptic complexes, D-loop formation and multiple invasions

As demonstrated above, RAD52 forms with RAD51 mixed and segmented filaments. We further tested the activity of theses filaments to pair with dsDNA and form SCs and D-loops (figure 6). Strikingly, we observed that the presence of RAD52 renders filaments 1,7 fold more active in contacting dsDNA homologous donors, with a great increase in SCs formation (from 43 to 72 %), demonstrating that mixed filaments are more efficient in establishing contacts with dsDNA donor (figure 6 A, C). This positive effect on SC formation was associated with a significant increase in stable D-loop intermediate formation from 4,9 to 13,8 % (in the absence and in the presence of RAD52 respectively, figure 6 D, E). It was partially independent of homology since the proportion of joint-molecules formed between mixed RAD51-RAD52 filament and a dsDNA heterologous donor is also significantly enhanced comparing to pairings involving pure RAD51 filaments and heterologous donors (figure 6 C). Increasing the RAD52 concentration in the reaction did not increase the representativeness of RAD52 within the filaments (as stated above) nor the quantity of SCs and D-loops but rather lead to an aggregation of dsDNA molecules with filaments, highlighting the high potential of RAD52 oligomer to interact with multiple DNA molecules bridging them together generating aggregates. In contrast, the addition of BRCA2 to the reaction had no significant effect on the proportion of joint molecules quantified either by TEM or in the D-loop assay although a slight (but not significant) increase in D-loop intermediates was reproductively observed, which may be explained by the fact that filaments formed in the presence of BRCA2 are longer and more stable (figure 6 C-E). Considering the architecture of SCs formed by the pairing of mixed RAD51-RAD52 filaments with homologous dsDNA donor, we did not show any change in the synaptic contact length. However, we noticed the frequent localization of RAD52 complexes on both sides of the junction zone, as if the presence of RAD52 was delineating the boundaries of the synapse. RAD52 was not detected along the junction zone. This result was confirmed by the specific labeling of RAD52 using an immunodetection assay (figure 6 A, B, F). The results are compatible with a first oligomer of RAD52 initiating contacts with the donor dsDNA and leading to an extension of the synaptic area to the next oligomer when homology is found. RAD52 could also play a role in delineating and then restricting the synaptic contact zone.

**figure 6:**
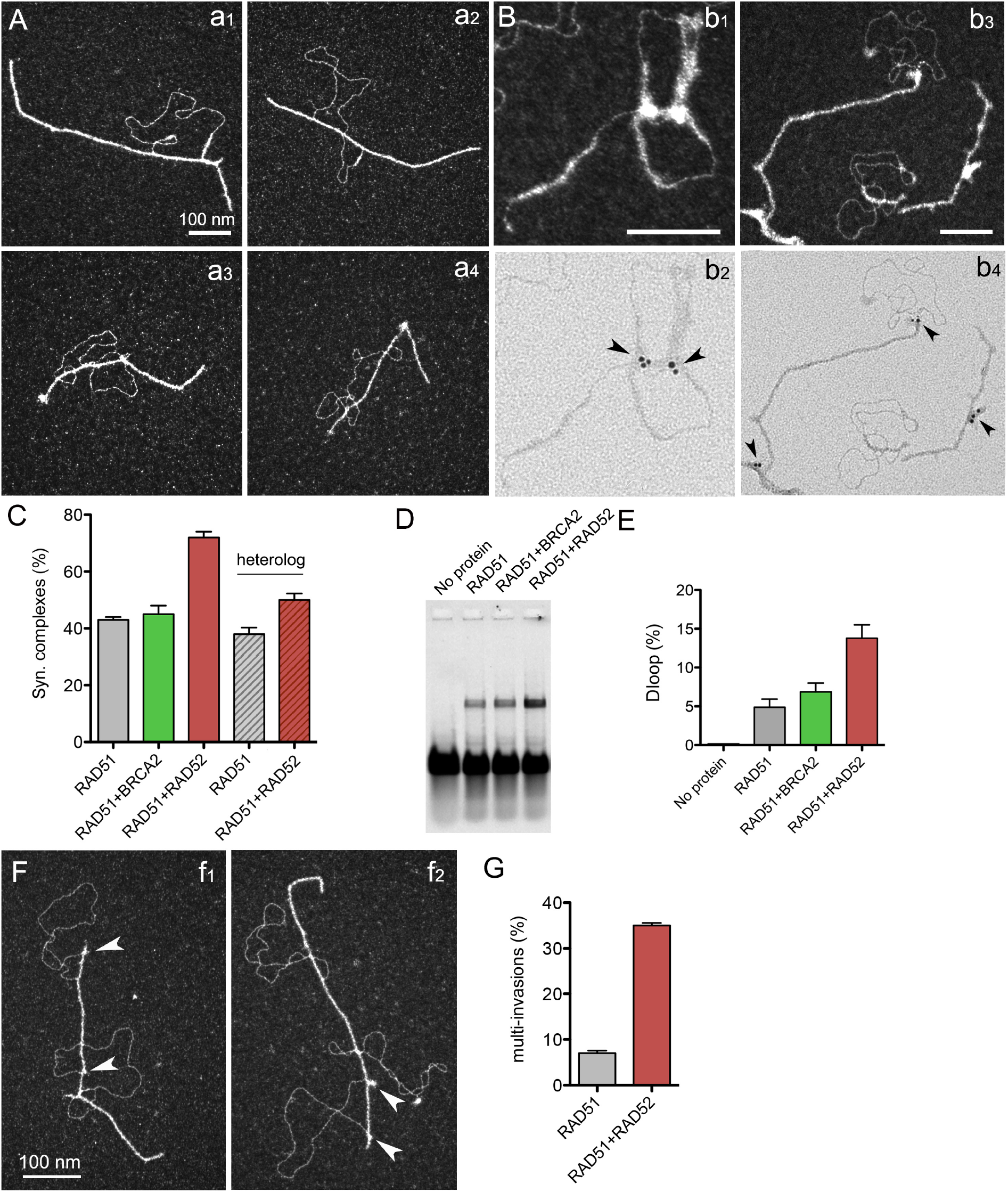
Mixed filaments formed in the presence of RAD52 are more active in contacting dsDNA and forming synaptic complexes, D-loops and multi-invasions. A. TEM images of joint molecules formed in the D-loop reaction by the pairing of mixed RAD51-RAD52 filaments with homologous dsDNA donor. B. Immunolabelling of the previous reaction using specific anti-RAD52 antibodies. C. Quantification of synaptic complexes formed during the D-loop reaction in presence of BRCA2 and RAD52 and also in presence of non homologous (heterologous) dsDNA. D. Deproteinized D-loop assay run on a 1% agarose gel and quantification of the D-loop yield (E). F. TEM images of multi-invasions identified in the D-loop reaction in presence of RAD52 and their qunatification (G).

Finally, we identified an interesting elevated rate of multi-invasion events within the population of RAD51-RAD52 mediated joint molecules, involving contacts with 2 or 3 donor DNAs (figure 6 F, G). Indeed, 35 % multi-invasions were quantified in conditions where RAD52 was present against 7 % in its absence. Again, bright discrete complexes were detected at the edges of the synaptic zone in these multi-invasion events. This result suggests that mixed RAD51-RAD52 filaments are not only active in their ability to contact dsDNA and form joint molecules but also in establishing multiple synapses by contacting several homologs at once. RAD52 could be involved in the alternative multi-invasion-induced rearrangement alternative mechanism described by Heyer et al. [75] (see discussion).

## Discussion

In this study, we aimed to characterize and understand how RAD52 can influence the formation and dynamics of the RAD51 nucleofilament in the presence of the well-defined RAD51 mediator BRCA2. Well conserved between unicellular eukaryotes like yeast *S. cerevisiae* and humans, the RAD52 functions seem to have evolved from its key role as main mediator of Rad51 filament in yeasts to a yet mysterious secondary role, away of its mediator functions, in mammals and vertebrates. Nonetheless, its synthetic lethality with BRCA2 pinpoints a yet critical role during HR that has proven difficult to uncover. We have approached the issue by using a combination of biochemistry and high resolution imaging by TEM, allowing us to visualize and analyze DNA-protein intermediates formed in reactions aiming to reconstitute *in vitro* the initial steps of HR, after the resection of the broken DNA ends. Our experiments allowed testing the putative collaboration of both BRCA2 and RAD52 in the substitution of RPA coated ssDNA to give rise to the nucleation and growth of a RAD51 nucleofilament. We confirmed in our analysis the spectacular ability of BRCA2 to promote RAD51 nucleation on ssDNA at sub-stoichiometric concentration (Figure 3), thus strongly indicating its role as the key RAD51 mediator in humans. We were also able to show how RAD52 tightly interacts and is able to lead RPA-coated ssDNA into a higher degree of compaction (Figure 2), thus partially inhibiting or limiting the RAD51 nucleation on the ssDNA section of the resected DNA ends (Figure 3 C, D). This result not only demonstrates that RAD52 is not a RAD51 mediator in humans, but quite the opposite it prevents RAD51 loading on RPA-ssDNA and helps limiting the length and continuity of the nucleofilaments it forms. Indeed, our data highlights the existence of two types of nucleofilaments depending on the presence and stoichiometry of these RAD51 partners: long, regular and continuous filaments mediated by an unopposed BRCA2 activity and shorter and discontinuous filaments occurring in the presence of RAD52, interspersed by clusters of RAD52 likely in the form of oligomers, that we have defined as mixed RAD51-RAD52 filaments (Figure 4). Interestingly, these mixed filaments are more active in establishing contacts with a dsDNA donor thus creating synaptic intermediates and D-loop (Figure 6). Also importantly, these mixed-filaments appear to be more proficient in establishing simultaneous multi-invasions of segmented RAD51-filaments (figure 6 F, G). In light of these results, we propose a model in which the two partner proteins could act sequentially, (1) At first, individually, protecting the newly formed ssDNA, and (2) secondly, synergistically, by promoting an efficient RAD51 nucleation, filament growth and HR stimulation with a limited length and more flexible synaptic abilities (see model in figure 7).

**figure 7:**
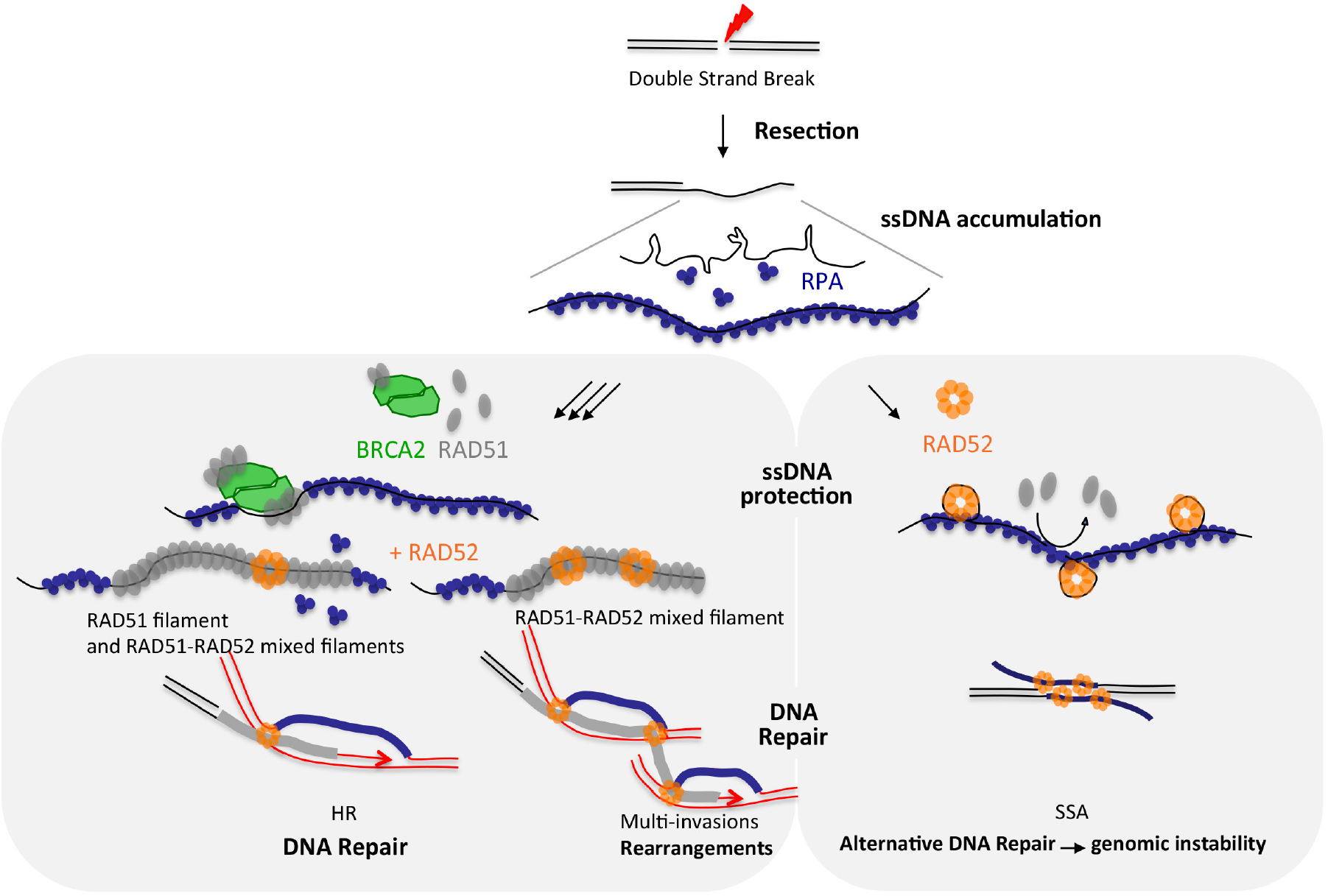
Model explaining the interplay between RAD52 and BRCA2 in HR early steps. Double strand break associated with resection lead to the accumulation of long ssDNA (overhang) that is rapidly covered by RPA protein. RAD52 can directly bind and compact RPA-ssDNA, inducing ssDNA protection and, in some context, associated with genomic instability. On the other hand, BRCA2 can bind and load RAD51 on RPA-ssDNA to promote filament assembly, ssDNA protection and HR. RAD52 can participate in the formation of some mixed and more active filaments to probe dsDNA donor and carry out homology search, but also more prone to multi-invasions and associated rearrangements.

### The ssDNA protection roles of BRCA2 and RAD52

The aforementioned protective functions of BRCA2 and RAD52 have been previously described, notably during replication stress [76] and would explain the synthetic lethality of the double mutants. Both proteins prevent extensive DNA degradation that may lead to cell death likely by binding to RPA-covered ssDNA and physically preventing the loading of nucleases onto the ssDNA. While on the one hand BRCA2 has an unparalleled ability to promote the recruitment and growth of RAD51 filaments (fully covering ssDNA), on the other hand RAD52 has a unique ability to freeze RPA-coated ssDNA into a more compact conformation, probably by wrapping the ssDNA around its oligomeric rings in the presence of RPA, with a preferential localization at the ss-dsDNA junction. Either one or the other presence may ensure the protection of ssDNA ends and thus function as redundant guardians of the genome. Accordingly, it was previously shown that under replication stress RAD52 associates with the nascent ssDNA, and the abrogation of the RAD52-ssDNA binding results in extensive nascent strand degradation by MRE11. It was then proposed that after replication fork stalling, RAD52 could be recruited onto ssDNA at perturbed forks, maintaining the fork in a closed conformation and thus playing a gatekeeper role [76]. We also observed *in vitro* that early binding of RAD52 to ssDNA inhibits RAD51 loading and filament installation likely by using the very same RPA-ssDNA compaction/gatekeeper properties. RAD52 is known to be the central player in the alternative and non-conservative DSB repair pathway of Single-Strand Annealing (SSA) that seal a break exposing two long complementary ssDNA sequences revealed after resection in a RAD51-independent manner [77,78]. Thus, in certain contexts, it might be possible that a precocious or excessive binding of RAD52 to the ssDNA-RPA overhangs may lead to SSA instead of HR if significant homologies are exposed, a deletion prone outcome putting at risk for genome integrity. It has been shown that this same SSA pathway is activated in HR-deficient cells, particularly in BRCA2/1-deficient tumoral cells where absence of functional BRCA2/1 has been associated with mutational signatures and ‘BRCAness’ phenotype [79]. In this pathological context, SSA ensures cell survival at expense of increased genome instability, thus likely being responsible of the progression and survival of the tumour’s cells. Our data sheds some light on the way RAD52 may regulate the recruitment of RAD51 via BRCA2, and how in the absence of this RAD51 filament, its ability to wrap ssDNA, promote proximity of ssDNA segments and its annealing properties may help survive cells experiencing DNA DSBs and replication stress by unusual though effective pathways of alternative repair like SSA.

### Recombination associated roles of BRCA2 and RAD52

The recombinogenic function of BRCA2 and RAD52 is related to their cooperation with RAD51 for the formation and activity of the presynaptic nucleofilament. Here we show that the role of RAD52 in the HR pathway could be explained by its direct participation in mixed nucleofilaments, making them more active and flexible during the homology search. We have specifically identified discrete RAD52 complexes within RAD51 filaments that we propose are composed of RAD52 oligomers around which ssDNA is wrapped, thus explaining the decrease in length of mixed RAD51-RAD52 filaments compared to otherwise “pure” or RAD51-only filaments. We described in a previous work how the yeast *S. cerevisiae* Rad52 protein was also able to form mixed filaments with Rad51 that were more resistant to the Srs2 helicase antirecombinase activity, as it was also the case for the Rad55 and Rad57 Rad51-paralogs [40,80]. In yeast, we were able to determine that the more Rad52 discrete complexes within the filaments, the more resistant they were to Srs2-led dismantling, suggesting that their presence may physically block the access or progression of antirecombinases along the DNA. The presence of human RAD52 within RAD51 filaments could have the same protective function against yet to be fully identified antirecombinase activities that may promote filament dismantling. Rad52 will thus work again as a gatekeeper within the Rad51-filament, as it does in replication fork contexts.

### Particular properties of human RAD51 nucleofilaments

In this work we have been able to visualize by using TEM, human RAD51-mediated joint molecules and characterize their architecture. Our observations clearly indicate that contrary to what happens in other well-studied models, the human RAD51 filaments have a strong ability to pair with dsDNA donors and form SC by themselves without the involvement of any third partner. Indeed, an unexpectedly high proportion (43%) of RAD51 filaments appeared as dsDNA-paired. From these filaments, only a few proportion were undergoing strand alignment and displacement of its complementary strand to create a D-loop, a distinction that can only be identified thanks to the use of our molecular imaging approaches. In addition, we showed that this efficiency of human RAD51 to probe and establish contacts with dsDNA is mostly independent of the homology of the sequences. An important proportion of the SC (38%) can be established with heterologous dsDNA donors. However, an extension of the synaptic contact zone was detected in the SC formed with homology-containing DNA donors, suggesting a change of the synaptic architecture once homology is found. Recombinases have two distinct DNA binding sites: (i) The site I that binds to ssDNA to form the presynaptic filament and (ii) the site II involved in contacting the donor dsDNA to mediate homology search [81]. It was proposed that the presynaptic filament would make transient and non-sequence specific contacts with an incoming dsDNA via site II to form a nascent three-stranded SC [20,82]. Two flexible loops of site I, L1 and L2, have been shown to respectively participate in the contacts with the donor dsDNA to form a first three-stranded SC, and then, providing sufficient homology is found, allow the transition to a second three-stranded SC where base flipping and strand alignment has occurred (L2)[83]. We believe that we have identified here the 2 types of three-stranded SCs: SCs formed with the heterologous dsDNA with a very short contact zone maintained by site II and L1 of RAD51, and the second SCs formed with the homologous dsDNA where the bases of the dsDNA have flipped thanks to homology and L2 and are in front of the complementary bases of the ssDNA of the filament, with a large synaptic area still maintained by RAD51.

While phage and bacterial recombinases are able to form SCs and D-loops autonomously, eukaryotic Rad51 relies on other proteins to achieve greater complexity of HR regulation. We have previously shown that ScRad51 filaments absolutely require ScRad54 to pair with dsDNA and engage in a SC during homology search, ScRad54 acting as bridging factor of Rad51 to dsDNA in a function analogous to the DNA binding site II [72]. ScRad54 plays a second essential function by converting the SC to a D-loop through strand alignment and RAD51 removal [26,72]. What we showed in this study suggests that human RAD51 functions differently and can perform dsDNA probing to form SCs autonomously, but can be assisted by RAD52 and its bridging activity to facilitate this step, while we can expect RAD54 to take over and stimulate the transition of these SCs to D-loops. Another interesting biochemical feature of human RAD51 is its ability to polymerize and bind stably to dsDNA as efficiently as to ssDNA [84]. However RAD51 was shown to display faster association on ssDNA in correlation with higher ssDNA flexibility [85,86] but slower dissociation from dsDNA, the RAD51-dsDNA filament being stable once formed [87]. It has been proposed that The presence of RAD51 on dsDNA has been evidenced *in vivo* in certain contexts, especially in the absence of RAD54-like proteins that desassemble toxic dsDNA-bound RAD51 complexes and prevent accumulation of nonrepair-associated RAD51 foci, pointing here again to the cooperation of important factor in the regulation RAD51 [65,88–90]. in spite of the fact that only filaments assembled on ssDNA are functional for homology search and strand exchange, one could still imagine that the transient and ephemeral polymerization of RAD51 on dsDNA would play a role, especially in the remodeling of certain DNA-protein complexes.

### A role of RAD52 in multi-invasion ?

A rearrangement mechanism has been recently described, based on HR mediated simultaneous invasions of two intact donors by a unique broken DNA end. The multi-invasion (MI) byproducts have been shown to be associated with chromosomal translocations [75]. Here we show that by forming fragmented mixed filaments, RAD51 and RAD52 are more likely to promote *in vitro* multi-invasions of distinct homologous dsDNA. In addition, in MI-synaptic complexes, RAD52 borders the synaptic zones. We propose that RAD52 may play a similar role in humans by enabling stable multiple invasions. These properties may be of special importance in the context of highly repetitive genome regions, wher multi-invasions may lead to cycles of deletion or expansion of repeated sequences.

## Acknowledgements

We thank the members of our laboratories for their helpfull comments and suggestions. This work was supported by grants from ANR (FIRE 17-CE12-0015), Paris-Sud University (MRM), CIHR FDN-388879 to J.Y.M. J.Y.M. is a Canada Research Chair in DNA repair and Cancer Therapeutics.

## Author contribution

A.A.M. and P.D. performed in vitro TEM and biochemical experiments. T.P. producted and purified RAD52 and BRCA2 proteins, and X.V. and X.B. producted and purified RAD51 and RPA. J.G.B. performed Homologous recombination in vivo assay. P.D., E.L.C., G.M.B and J.Y.M. designed the study and analyzed the results. P.D. wrote the manuscript with the help of G.M.B., E.L.C and J.Y.M.

## Conflict of interest statement

The authors declare no conflicts of interest.

## Notes

### Competing Interest Statement

The authors have declared no competing interest.

