## supplementary figures for "Human RAD52 stimulates the RAD51-mediated homology search"

**Supplemental data.**

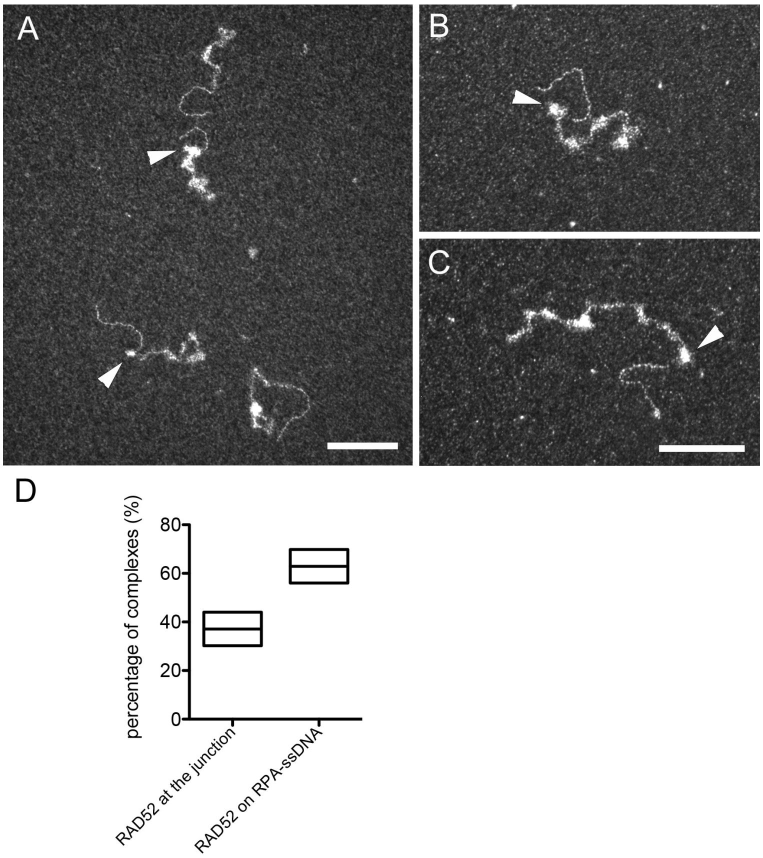


<u>**Supplementary figure S1:** RAD52 preferentially binds to the ss-ds junction.</u>
The ss-ds DNA substrate is first incubated with saturated amount of RPA then 0,2 μM RAD52 are added to the reaction. (A-C) TEM analysis of the RAD52-RPA-DNA complexes. RAD52 forms discrete complexes with RPA-ssDNA. (D) 39 % of RAD52-RPA-DNA complexes are quantified with a specific binding of RAD52 to the ss-ds junction. The scale represents 100 nm.

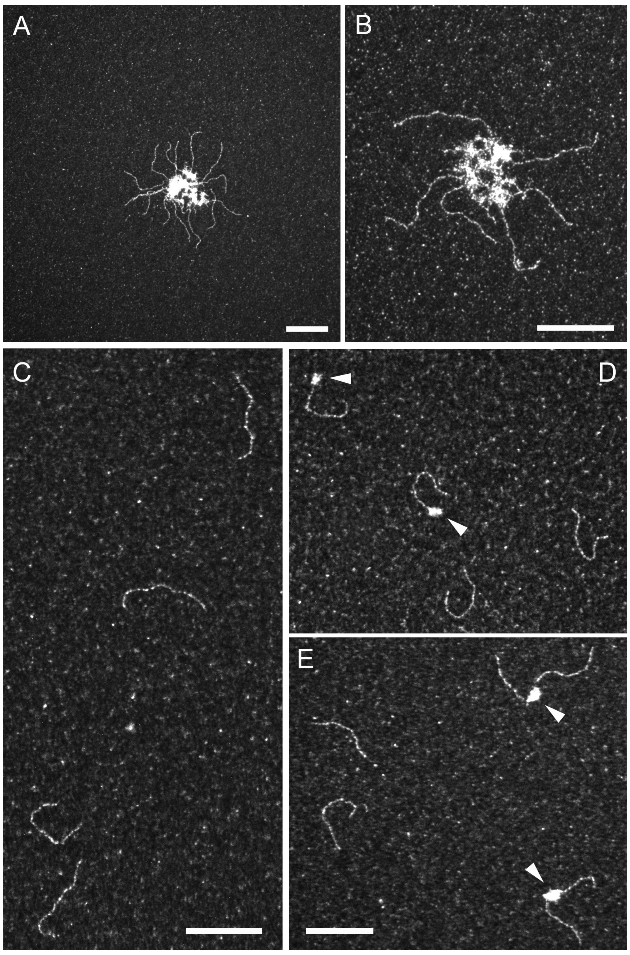


<u>**Supplementary figure S2:** RAD52 binding to different DNA substrates in absence of RPA.</u>

(A-B) ss-dsDNA substrate containing a 400 bp dsDNA extended with a long 1040 nt single stranded overhang is incubated with 0,2 μM of RAD52. RAD52 induces the bridging and aggregation of the ssDNA parts. (C) A blunt 400 bp dsDNA is incubated with 0,2 μM of RAD52. No binding to dsDNA or agregation are detected in these conditions. (D-E) An ss-dsDNA substrate containing a 400 bp dsDNA extended with a short 4 nt single stranded overhang is incubated with 0,2 μM of RAD52. 4 nt are sufficient to allow the recruitment of RAD52 and the formation of bridges between 2 or 3 DNA substrates. The scale bar represents 100 nm.

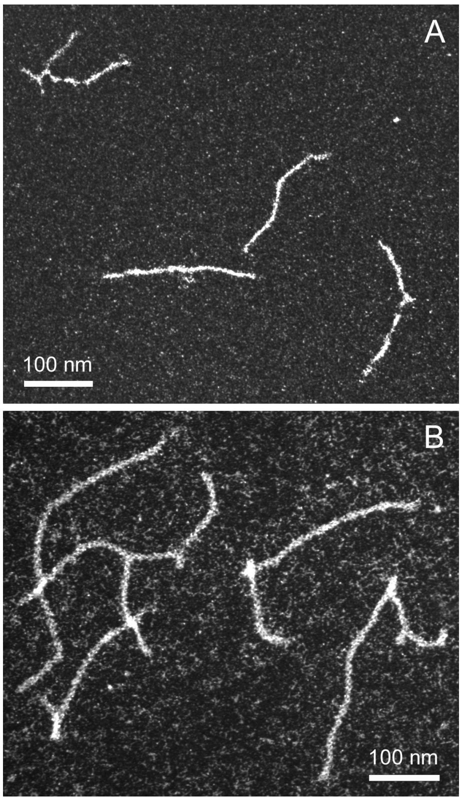


**Supplementary figure S3:** RAD51 behaviour on ss-dsDNA substrate

(A) 9 μM of ss-dsDNA substrate containing a 400 bp dsDNA extended with a long 1040 nt single stranded overhang is incubated with 3 μM of RAD51, in absence of RPA. Filaments are shorter than those formed in presence of RPA added juste after RAD51. RAD51 similarly bind to ss and dsDNA. (B) Filaments are assembled on same substrate in presence of 1 μM of RAD52. Mixed filaments are formed with the formation of discrete complexes. The background reflected the covering of TEM grids with excess of proteins.
